# ShinyMultiome.UiO: An interactive open-source framework utilizing Seurat Objects for visualizing single-cell Multiomes

**DOI:** 10.1101/2023.06.20.545756

**Authors:** Akshay Akshay, Ankush Sharma, Ragnhild Eskeland

## Abstract

**Motivation:** Single-cell genomics has been revolutionized by the advent of single-cell Multiome sequencing which allows for simultaneous profiling of chromatin accessibility and gene expression in individual nuclei. The analysis and interpretation of large and often complex scMultiome datasets requires in-depth knowledge of computational programming.

**Results:** We present ShinyMultiome.UiO, a user-friendly, integrative, and opensource web-based tool that supports Seurat objects for visualization of single-cell Multiome. The ShinyMultiome.UiO facilitates interactive reporting and comprehensive characterization of cellular heterogeneity and regulatory landscapes. With ShinyMultiome.UiO, users can facilitate collaborative efforts for interpretation of single-cell Multiome data and make it available to the public.

**Availability and implementation:** https://github.com/EskelandLab/ShinyMultiomeUiO and demo server, https://cancell.medisin.uio.no/ShinyMultiome.UiO/ set up with a PBMC reference dataset.

**Contact:** &

## 1. Introduction

Single-cell multi-omics is emerging as a powerful approach for understanding biological processes and unravelling cellular heterogeneity in development, tissues, and disease. We previously developed ShinyArchR.UiO, an open-access web application built using the R programming language and the Shiny framework, which was designed primarily for the analysis of single-cell Assay for Transposase-Accessible Chromatin sequencing (scATAC-seq) by ArchR (Sharma *et al*., 2021; Granja *et al*., 2021). Additionally, ShinyArchR.UiO can plot multi-omics integration of single-cell RNA sequencing (scRNA-seq) and scATAC-seq, and can be used in conjunction with ShinyCell to report and share data from the two individually collected modalities (Ouyang *et al*., 2021; Sharma *et al*., 2021). Single-cell Multiome (scMultiome) combines scATAC-seq with scRNA-seq from the same nuclei, and the direct link between the two modalities have shed new light on gene regulatory mechanisms in distinct cellular states (Belhocine *et al*., 2021; Ma *et al*., 2020). However, to fully exploit scMultiome data, sophisticated computa-tional methods and userfriendly tools are re-quired. The increase in intra-modality Seurat annotation for scMultiome data calls for a web-based interactive reporting tool to enhance Findable, Accessible, Interoperable and Reusable data sharing.

## 2. An Overview of ShinyMultiome.UiO

ShinyMultiome.UiO features an interactive and intuitive web-based interface for reporting and visualizing single-cell Multiome data from the same cell analysed using Seurat and Signac integration method (Stuart *et al*., 2019, 2021; Satija *et al*., 2015). We demonstrate ShinyMultiome.UiO on our local server with a public scMultiome dataset (v1.0) comprising 10K Human peripheral blood mononuclear cells (PBMCs) sequenced on ChromiumX (Fig. 1) (10x Genomics). Plot height and width can be customized before a high-quality tiff, png, or pdf is generated.

**Figure 1.**
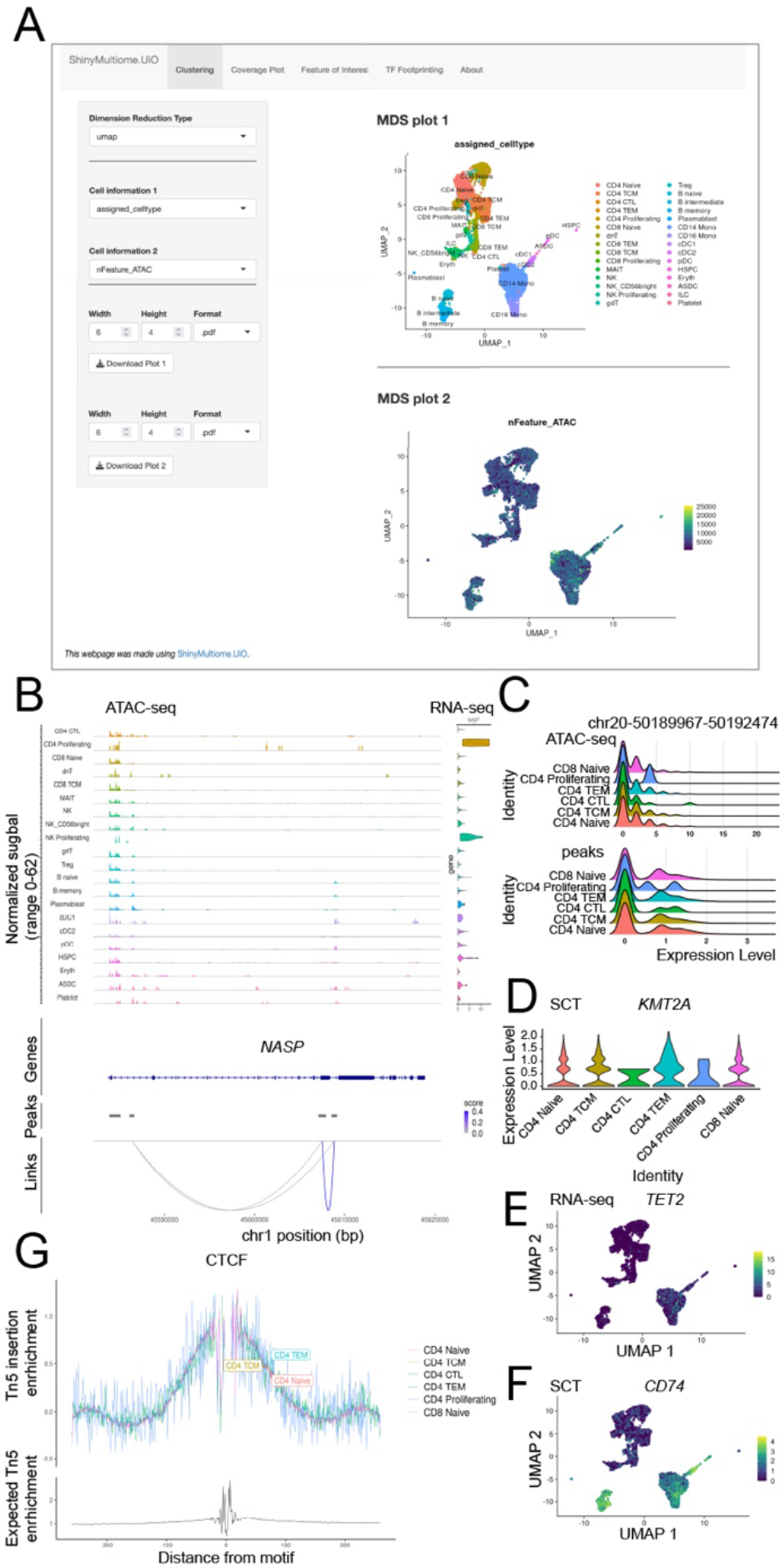
ShinyMultiome.UiO display four different web interfaces in tabs where the user can generate multiple plots from scMultiome data. The about tab enclose information about the tool and all references. (a) Clustering is the front user-interface and allow for selection and exploration of three different dimension reduction types (PCA, LSI and UMAPs) in two sizeable plots, displayed top and bottom, as demonstrated with clusters from a scMultiome dataset of PBMCs and nFeatureATAC. Multiple cell information can be selected and saved as tiff, png or pdf. (b) The Coverage Plot tab shows peaks for scATAC-seq modality on different clusters with adjacent violin plots representing the gene of interest expression level in PBMC clusters on scRNA-seq modality. In the Feature of interest tab, two plots can be viewed simultaneously of different feature comparisons using ridge plot, violin plot or UMAPs. Different cellular clusters can manually be added or removed in the Cell Types panel when using the ridge plot and violin plot. (c) Two representative ridge plots visualizing ATAC signals and peaks across the *CEBPB* locus in six selected PBMCs clusters on Genome build GRCh38/hg38. (d) An example of a violin plot of gene expression of *KMT2A* in SCT assay across selected PBMC clusters. (e) A UMAP plot of *TET2* gene expression in RNA-seq assay and (f) a UMAP plot showing gene expression of *CD74* on a SCT assay. (g) TF Foot-printing of CTCF in six selected PBMC clusters.

## 3. Key features of ShinyMultiome.UiO

ShinyMultiome.UiO can seamlessly integrate single-cell scMultiome data, enabling users to explore the relationship between chromatin accessibility and gene expression in a unified framework. This integration allows for comprehensive analysis of cell populations, identification of cell-type specific regulatory regions, and characterization of gene regulatory networks. ShinyMultiome.UiO offers the following key features:

1. Joint visualization of chromatin accessibility and gene expression: ShinyMultiome.UiO provides interactive plots and visualizations enabling users to identify co-regulated gene sets and potential regulatory elements.
2. Dimensionality reduction and clustering: ShinyMultiome.UiO supports popular dimensionality reduction techniques such as LSI, UMAP, and PCA, allowing users to visualize and cluster cells based on their multi-modal profiles.
3. Browser view of scATAC-seq peaks: Genomic coordinates or gene symbol can be selected for visualization of chromatin opening in user selected clusters with the option of extending the browser window upstream and downstream. Mapped peaks and links between chromatin opening and correlated genes are displayed. Cluster based gene expression is represented by violin plots.
4. Transcription factor footprinting: Users can rapidly plot motif footprint of TF regulators within clusters of choice accounting for ex-pected Tn5 enrichment.

## 4. Basic usage

ShinyMultiome.UiO is an R based tool that can be set up on different operating systems. This process can be executed in two simple steps:

i. Download or git clone https://github.com/EskelandLab/Shiny-MultiomeUiO
ii. Provide a path to Seurat objects (Stuart and Srivastava) in RDS format and fragment file in *global*.*R*. The initiation of the shiny app takes few minutes on a quad-core 8 gigabytes of RAM of data representing more than 10000 cells.

## 5. Open source and data sharing

ShinyMultiome.UiO web-interface makes exploring and understanding of scMultiome data easy and can be run locally or made available on a locally hosted server. Alternatively, an open-source shiny-server can be set up (RStudio Team, 2020). ShinyMultiome.UiO is an opensource R-code tool for a broader range of users than experienced computational scientists. The source code for ShinyMultiome.UiO can easily be customised.

## Conclusions

The ShinyMultiome.UiO tool provides essential visualization options such as dimensionality reduction, chromatin accessibility and transcription factor footprints. With the ShinyMultiome.UiO, single-cell multiome data can be made available to a wider audience fostering sustainable research practices and reducing the carbon footprint associated with repetitive data analysis.

## Author Contributions and Funding

AA and AS developed the code with scientific input and tests performed by RE. RE and AS wrote the manuscript with input from AA. All authors have read and approved the final manuscript. This work was partly supported by the Research Council of Norway through its Centres of Excellence funding scheme, project number 262652. Conflict of interest, none declared. Disclaimer – AS is an employee of Acerta-Pharma, The Netherlands.

## Acknowledgments

We would like to acknowledge Knut Waagan at the Department for Research Computing, UiO for help to set up the CanCell server and ShinyMultiome.UiO demo. We thank Mo-hamed Abdelhalim for testing ShinyMulti-ome.UiO on Microsoft Windows operating system.

## References

10x Genomics https://www.10xgenomics.com/.

Belhocine, K. et al. (2021) Single-Cell Multiomics: Simultaneous Epigenetic and Transcriptional Profiling. Genetic Engineering & Biotechnology News, 41, 66–68.

Granja, J.M. et al. (2021) ArchR is a scalable software package for integrative single-cell chromatin accessibility analysis. Nature Genetics, 53, 403–411.

Ma, S. et al. (2020) Chromatin Potential Identified by Shared Single-Cell Profiling of RNA and Chromatin. Cell, 183, 1103–1116.e20.

Ouyang, J.F. et al. (2021) ShinyCell: simple and shara-ble visualization of single-cell gene expression data. Bioinformatics, 37, 3374–3376.

RStudio Team (2020) RStudio: Integrated Development for R. RStudio, PBC, Boston, MA.

Satija, R. et al. (2015) Spatial reconstruction of single-cell gene expression data. Nat Biotechnol, 33, 495–502.

Sharma, A. et al. (2021) ShinyArchR.UiO: user-friendly, integrative and open-source tool for visualization of single-cell ATAC-seq data using ArchR. Bioinformatics.

Stuart, T. et al. (2019) Comprehensive Integration of Single-Cell Data. Cell, 177, 1888-1902.e21.

Stuart, T. et al. (2021) Single-cell chromatin state analysis with Signac. Nat Methods, 18, 1333– 1341.

Stuart, T. and Srivastava, A. Analysis of Single-Cell Chromatin Data. https://stuartlab.org/signac/.

